# COMADRE: a global database of animal demography

**DOI:** 10.1101/027821

**Authors:** Roberto Salguero-Gómez, Owen R. Jones, C. Ruth Archer, Christoph Bein, Hendrik de Buhr, Claudia Farack, Fränce Gottschalk, Alexander Hartmann, Anne Henning, Gabriel Hoppe, Gesa Römer, Tara Ruoff, Veronika Sommer, Julia Wille, Jakob Voigt, Stefan Zeh, Dirk Vieregg, Yvonne M. Buckley, Judy Che-Castaldo, David Hodgson, Alexander Scheuerlein, Hal Caswell, James W. Vaupel

## Abstract

1. The open-access scientific philosophy has been widely adopted and proven to promote considerable progress in the fields of ecology and evolution. Openaccess global databases now exist on animal migration, the distribution of species, and conservation status, to mention a few. However, a gap exists for databases on population dynamics spanning the rich diversity of the animal kingdom. This information is fundamental to our understanding of the conditions that have shaped variation in animal life histories and their relationships with the environment. Furthermore, an animal population’s schedules of reproduction and mortality determine its invasive potential, and its risk of local extinction, which are at the core of conservation biology.
2. Matrix population models (MPMs) are among the most widely used demographic tools by animal ecologists. MPMs project population dynamics in terms of reproduction, mortality, and development over the entire life cycle. The results of MPMs have direct biological interpretations, facilitating comparisons among animal species as different as *Caenorhabditis elegans, Loxodonta africana* and *Homo sapiens.*
3. Thousand of animal demographic records exist in the form of MPMs, but they are dispersed throughout the literature, rendering comparative analyses difficult. Here, we introduce the COMADRE Animal Matrix Database version 1.0.0, an open-source online repository containing data on 402 species worldwide, from 272 studies, with a total of 1,575 population projection matrices. COMADRE also contains ancillary information (e.g. ecoregion, taxonomy, biogeography, etc.) that facilitates interpretation of the numerous demographic metrics that can be derived from its MPMs.
4. *Synthesis:* We introduce the COMADRE Animal Matrix Database, a resource for animal demography. The open access nature of this database, together with its ancillary information will facilitate comparative analysis, as will the growing availability of databases focusing on other traits, and tools to query and combine them. Through future frequent updates of COMADRE, and its integration with other online resources, we encourage animal ecologists to tackle global ecological and evolutionary questions with unprecedented statistical power.

## Introduction

An understanding of the drivers and consequences of variation in reproduction and mortality throughout the life cycle is at the heart of population biology, evolution, ecology, and allied fields (Metcalf & Pavard 2006; Salguero-Gómez & de Kroon 2010; Salguero-Gómez *et al.* 2015). Although demography is essential to understand and predict population dynamics, no single repository integrates these data. This is mainly because most biological data sources are scattered and biological data types are heterogeneous (Hoffmann *et al.* 2014). Moreover, demographic data pose challenges for standardization due to the different formats and terminology (Lebreton 2012; Conde *et al.* unpublished). This makes it challenging to create a single demographic data repository across multiple species. However, there are important efforts towards compiling these data such as the Global Population Dynamics Database (GPDD, Inchausti & Halley 2001) and the Living Planet Index (LPI, Collen *et al.* 2009) holding population time-series data, BIDDABA (Lebreton *et al.* 2012), and the Primate Life History Database (PLHD, Strier *et al.* 2010) containing demographic information for birds and primates respectively. Although these examples have advanced the field of population biology, they are limited in either demographic detail (GPDD, LPI) or taxonomic scope (BIDDABA, PLHD, WBI).

A mechanistic understanding of how and why populations invade, grow, decline, or go locally extinct, requires data and methods that provide insights into age/size/ontogeny-based structure, such as Matrix Population Models (*MPMs* hereafter; Caswell 2001). MPMs have become the staple method describing the structured demography of animal populations. The widespread use of MPMs stems from their well-understood mathematical foundations and tractability (Caswell 2001), coupled with the clear biological interpretations of the analytical outputs (de Kroon *et al.* 1986; Silvertown, Franco & Menges 1996; de Kroon, van Groenendael & Ehrén 2000). Briefly, an MPM divides the life cycle into discrete stages and projects the population through time in terms of probabilities of survival and of transitions among stages, and of the contributions to sexual or clonal reproduction at each stage. The stages of the life cycle can be chosen based on a compromise between the biology of the species and the availability of data, and the projection interval can vary from days (e.g. Buston & García 2007) to years (e.g. Edmunds *et al.* 2015), depending on the species and question.

As is the case with plants (Salguero-Gómez *et al.* 2015), a large number of MPMs have been published on species in the animal kingdom since the models were introduced in the 1940s (Bernadelli 1941; Leslie 1945) (Figure 1). Underlining the general utility of MPMs, these models have been used to address diverse topics including conservation biology (e.g., Crouse, Crowder & Caswell 1987; Jenouvrier *et al.* 2012), evolutionary biology (e.g., Kawecki 1995), ecotoxicology (e.g., Charles *et al.* 2009), invasion biology (e.g., Neubert & Parker 2004), and resource management (e.g., Salomon *et al.* 2013). MPMs have been employed to study species as taxonomically distinct as *Caenorhabditis elegans, Loxodonta africana* and *Homo sapiens,* and in geographically diverse regions with studies in every major biome (Figure 2.A & 2.B).

**Figure 1.**
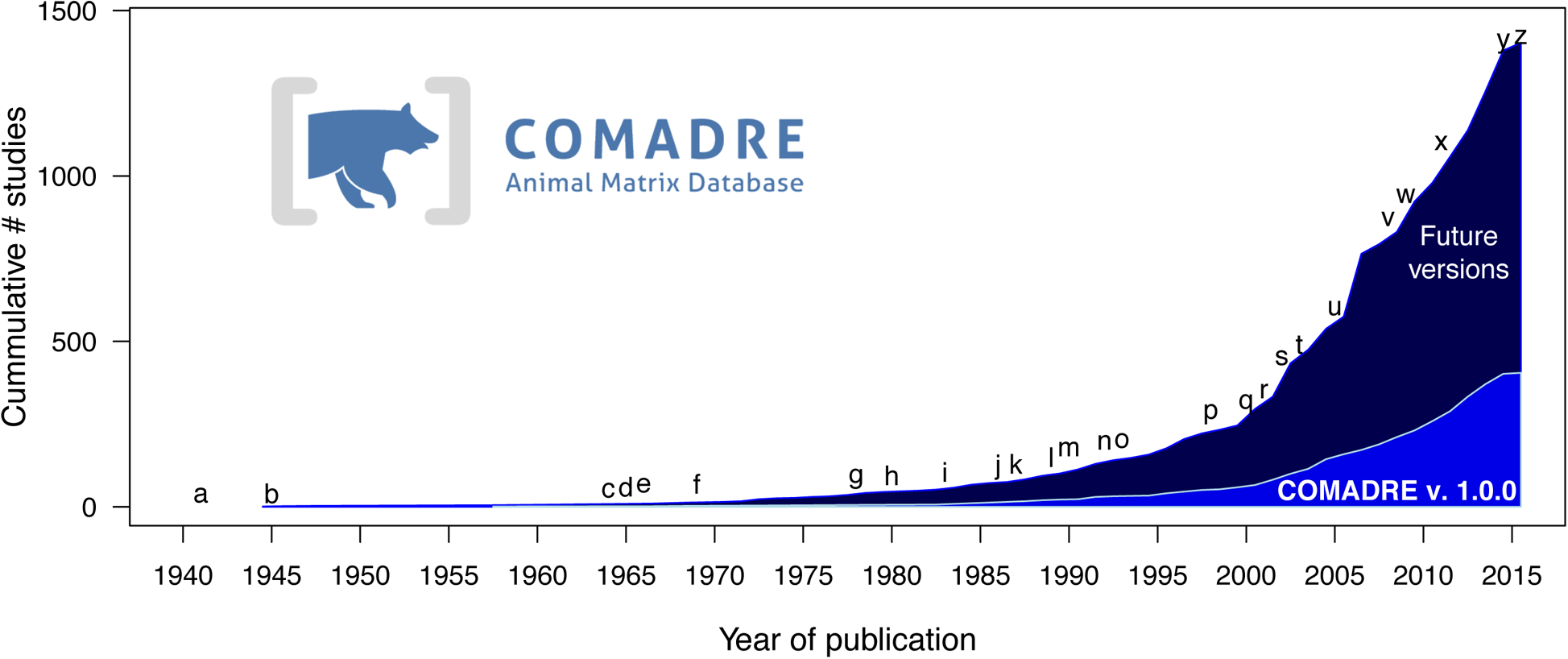
Time-line of the cumulative number of studies published up until May 31^st^ 2015 containing matrix population models (MPMs) of animals in peer-reviewed journals, books, reports and theses. The light blue background corresponds to studies released in COMADRE version 1.0.0; Dark blue corresponds to studies currently under inspection, to be incorporated in future versions. Pivotal events in the development of the COMADRE Animal Matrix Database: (*a*, *b*) first applications of matrix models in demography (Bernardelli 1941; Leslie 1945), (*c*) first application of MPMs to human demographic projections (Keyfitz 1964), (*d*) introduction of theory for stage-classified MPMs (Lefkovitch 1965), (*e*) first spatial MPM (Rogers 1966), (*f*) first nonlinear, density dependent MPMs for animal populations (Pennycuick 1969; Rabinovitch 1969), (*g*) first sensitivity analysis for stage-classified MPMs and calculation of selection gradients for animals (Caswell 1978), (*h*) first bifurcation analysis of density-dependent MPMs in animals (Levin & Goodyear 1980), (*i*) first calculation of the stochastic growth rate from an animal MPM (Cohen, Christensen & Goodyear 1983), (*j*) formalization of elasticity analyses for MPMs (de Kroon *et al.* 1986), (*k*) first application of elasticity analysis in conservation biology for *Caretta caretta* (Crouse, Crowder & Caswell 1987) and first Life Table Response Experiment analysis (Levin *et al.* 1987), (*l*) first edition of the *Matrix Population Models: Construction, Analysis, and Interpretation* (Caswell 1989), (*m*) publication of *Population Dynamics in Variable Environments* (Tuljapurkar 1990), (*n*) first presentation of multi-state mark-recapture methods for estimating stage-structured MPMs in animals (Nichols *et al.* 1992), (*o*) development of the first MPM from photo identification data (Brault & Caswell 1993), (*p*) one of the first studies to detail uncertainty in MPMs (Caswell *et al.* 1998), (*q*) first special feature on MPMs (Heppell, Pfister & de Kroon 2000), (*r*) 2^nd^ edition of *Matrix Population Models* (Caswell 2001), (s) publication of *Quantitative Conservation Biology: Theory and Practice of Population Viability Analysis* (Morris & Doak 2002) summarizing and stimulating applications of MPMs to conservation, (*t*) first application of matrix integro-difference equations to examine invasion speeds in animal populations (Caswell, Lensink & Neubert 2003), (*u*) first investigation of nonequillibrium properties, such as reactivity, for MPMs (Caswell & Neubert 2005), (*v*) first complete perturbation analysis for nonlinear animal MPMs (Caswell 2008), (*w*) introduction of individual stochasticity analyses for animal MPMs (Caswell 2009; Tuljapurkar *et al.* 2009), (*x*) foundation of the COMADRE database at the Max Planck Institute for Demographic Research, (*y*) release of the COMPADRE Plant Population Database 3.0 in www.compadre-db.org (Sept 11^th^ 2014), and (*z*) online release of the COMADRE Animal Matrix Database version 1.0.0 in www.comadre-db.org.

**Figure 2.**
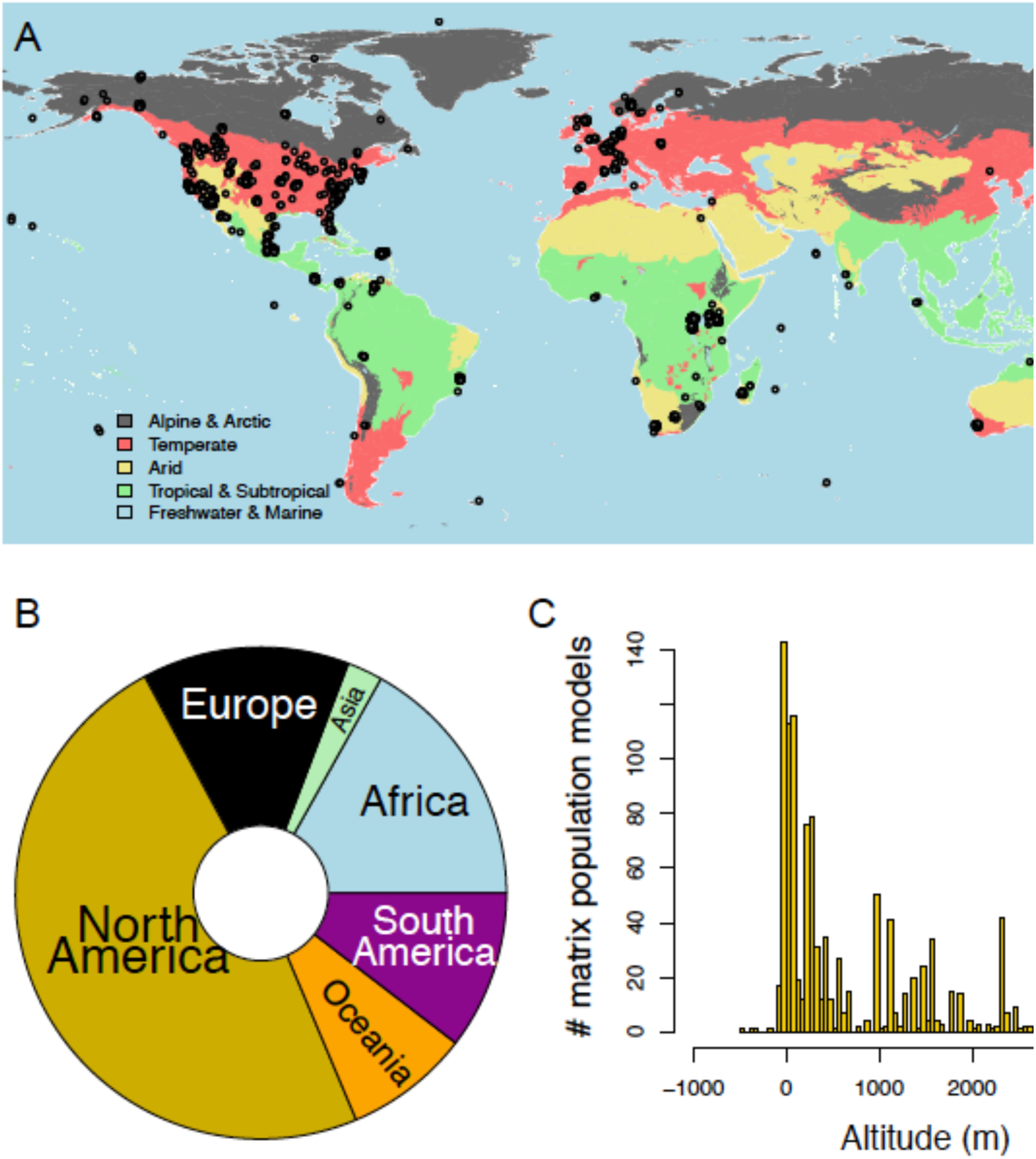
Geographic representation of animal demographic studies in COMADRE. **A**. Worldmap of studies in COMADRE 1.0.0. The points represent studied sites, and have been jittered to highlight temporal replication and close spatial overlap of certain studies. World map shows major habitats as color-coded background (See legend). **B**. Breakdown of studies by continent. **C.** Frequency of MPMs by altitude. Negative values typically indicate marine and freshwater sites.

Despite the growing availability of published MPMs and the fact that such models are inherently comparable, there have been few attempts to use MPMs in comparative analyses. Notable exceptions are the work by Sæther and Backer (2000) on birds, and Heppell, Caswell and Crowder (2000) on mammals, Vélez-Espino, Fox and McLaughlin (2006) on fish, and van de Kerk *et al.* (2013) on order Carnivora. These works illustrate the power of comparative approaches for robust generalizations by relating demographic estimates made from MPMs to interactions with the environment that form the basis for the evolution of life histories. One reason for the lack of comparative studies has historically been the paucity of readily available demographic data, compared to genetic data (e.g. Benson *et al.* 2013). This changed earlier this year, when Salguero-Gómez and colleagues (2015) released a database on plant demography, COMPADRE. Since its foundation in 1990, COMPADRE has prompted over 35 comparative plant demography studies ranging from senescence (Silvertown, Franco & Perez-Ishiwara 2001), to short-term population dynamics (Stott, Townley & Hodgson 2011), to the link between functional traits and demography (Adler *et al.* 2014). Here, we announce the release of COMPADRE’s sister database, COMADRE, which contains MPMs and associated metadata from the animal kingdom.

The main objectives of the COMADRE team are (i) to find, digitize, and systematically error-check published animal MPMs and supplement them with additional information (Table 1), (ii) to offer such information on an open access basis, and (iii) to develop *R* scripts to facilitate comparative analyses. The data described here are available at www.comadre-db.org. In this paper, we briefly describe COMADRE, and highlight the major differences and similarities between COMADRE and the sister-database on plants COMPADRE. In addition, we briefly report some geographic, taxonomic and modelling biases inherent in the database. Finally, we detail our vision for how COMADRE will expand and develop in the future, linking to other already existing open-access databases to address timely questions in animal ecology and evolution.

**Table 1.**
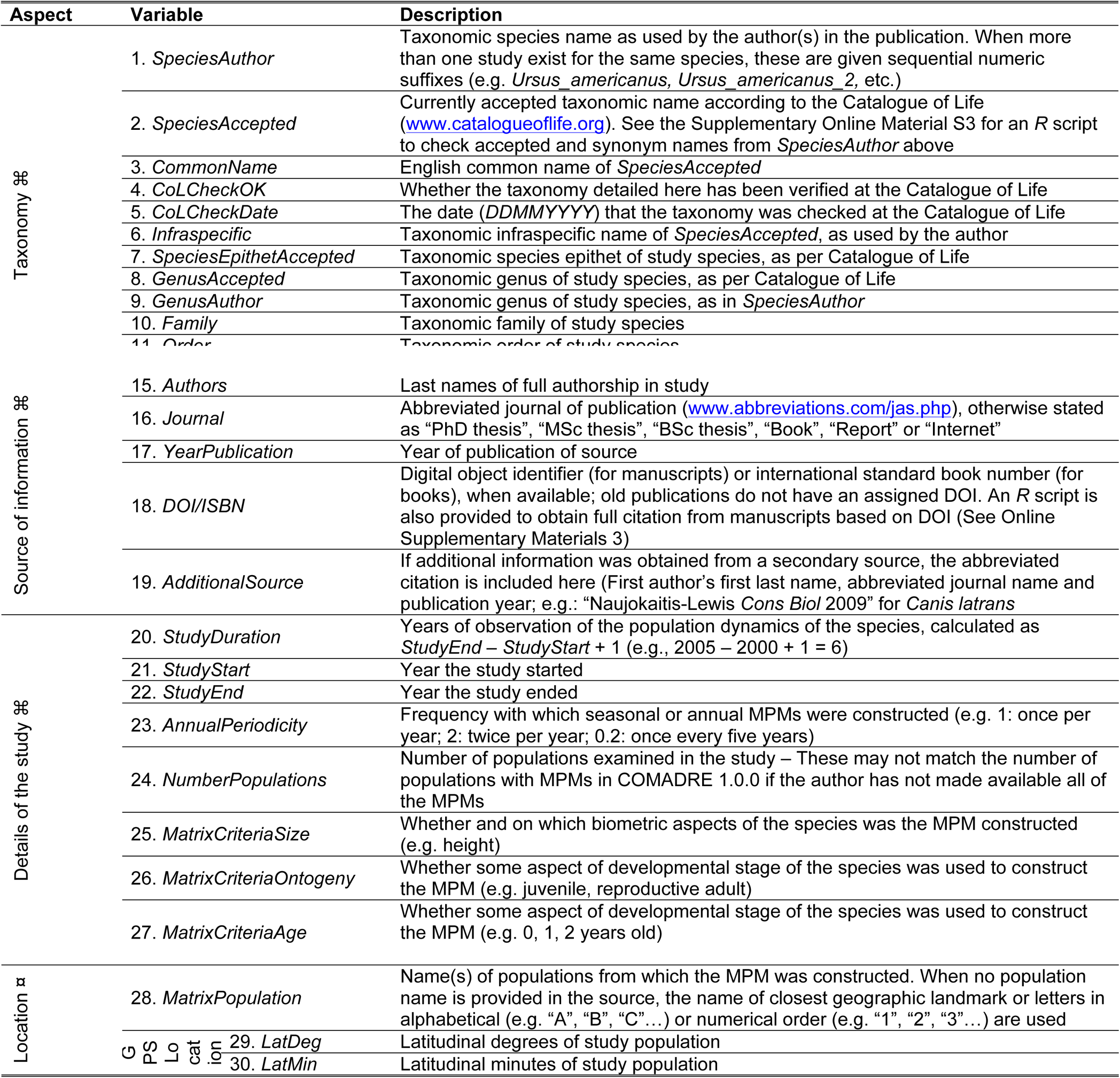
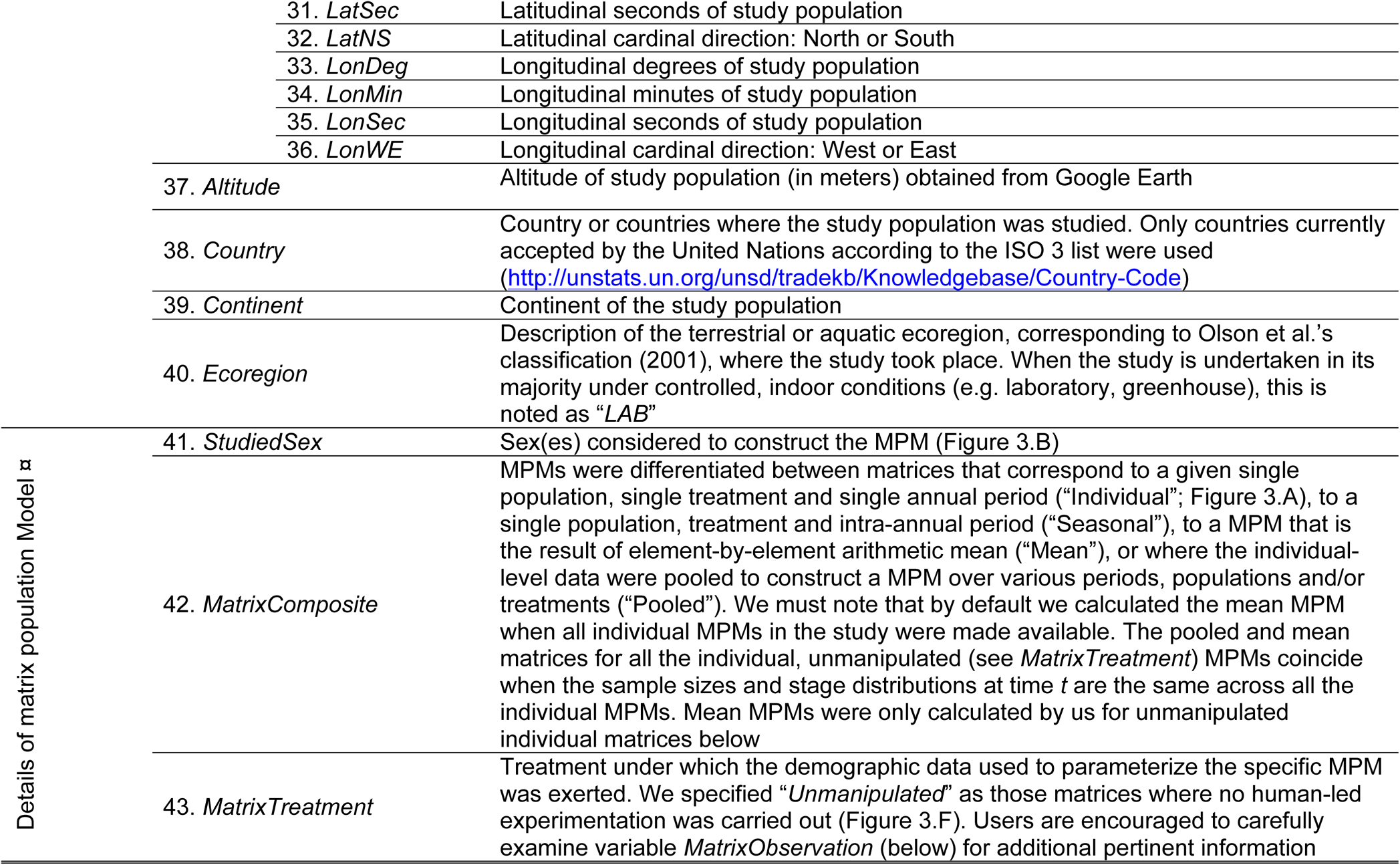
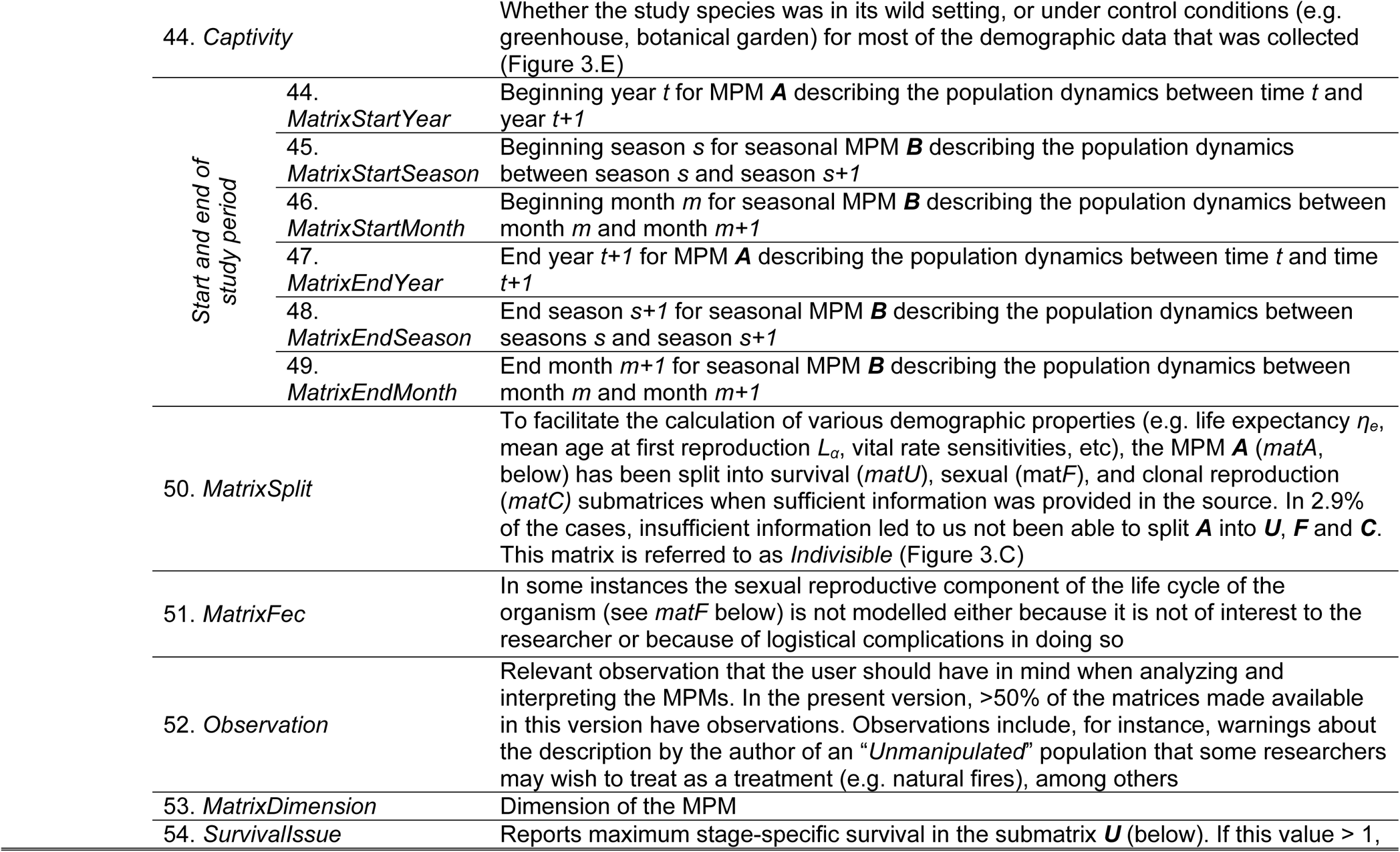
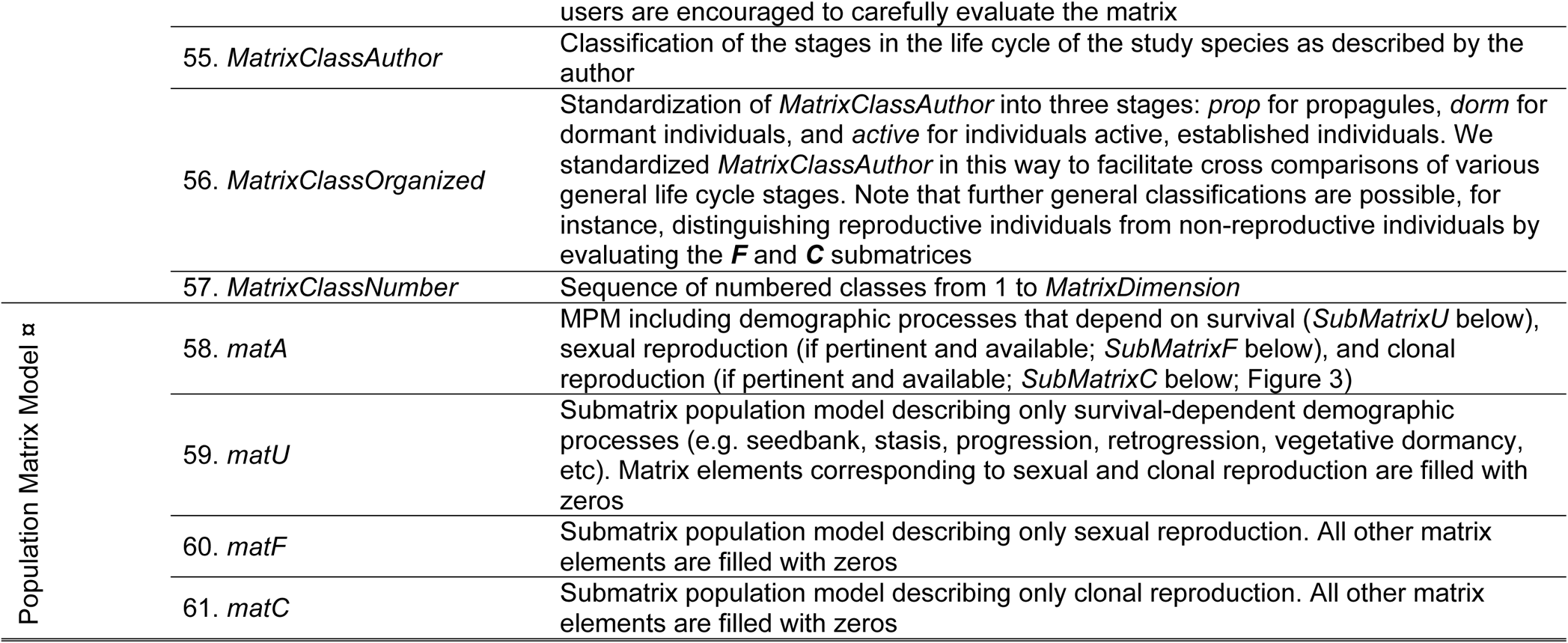
Variables in the COMADRE Animal Matrix Database, organized by six general aspects: taxonomy, source, details of study, geolocation, and Matrix Population Model (MPM). A more detailed description can be found in the user protocol of COMADRE at www.comadre-db.org. ⌘ implies information that is study-specific; n information that is MPM-specific. Variables 1-54 are archived in `comadre$metadata,` variables 55-57 in `comadre$matrixClass,` and variables 58-61 in `comadre$mat` in the COMADRE *R* data object open access available in the COMADRE online portal.

## COMADRE 1.0.0: A historical perspective

The accumulated number of publications reporting MPMs for animals has increased dramatically since MPMs were introduced in the 1940s (Figure 1). Important contributions to this history come from the introduction of new *types* of MPMs, and new *methods* for analysing them.

Matrix population models were largely ignored for twenty years after the work of Leslie (1945). This is partly because Leslie had also helped introduce life table calculations of population growth rate into ecology, and those methods were more computationally feasible in the days before computers (Caswell 2001)^1^.

The rediscovery of MPMs in the 1960s can be credited to three papers (Keyfitz 1964; Lefkovitch 1965; Rogers 1966). All of these papers focused on animals (yes, humans *are* animals). Keyfitz (1964) presented MPMs as tools for projecting population growth; his book (Keyfitz 1968) influenced a generation of animal ecologists. The first presentations of MPMs had assumed that age was the only i-state variable. Lefkovitch (1965), based on studies of laboratory populations of stored product insect pests, explicitly proposed stage-classified models based on the stages of the insect life cycle. Rogers (1966) introduced spatial, or multiregional, models for human populations, classifying individuals by age and spatial location, and modelling mortality, fertility, and migration between locations.

Other types of MPMs were introduced in the following years. The first seasonal, periodic MPM appeared in 1964 (Darwin & Williams 1964) in a study of seasonal harvesting as a control strategy for rabbits. The first density-dependent models appeared in 1969; Pennycuick *et al.* (1969) analyzed a population of great tits *(Parus major),* based on field data. Rabinovich (1969) comparing several density-dependent models, including a MPM to analyze laboratory populations of a parasitoid wasp. The first stochastic model for an animal population was the analysis by Cohen, Christensen and Goodyear (1983) of recruitment fluctuations in striped bass *(Morone saxatilis).* Invasion models, using matrix integrodifference equations, were first applied to bird populations by Caswell, Lensink and Neubert (2003).

Analytical methods have developed in parallel with their applications to animal populations. Many of these are listed in Figure 1. Some of these developments have provided new ways of constructing models (photo-identification methods, mark-recapture methods, vec-permutation matrix methods). Others have provided ways to extract additional information from the resulting MPM (sensitivity and elasticity analyses; LTRE decomposition analyses; stability and bifurcation analyses for nonlinear models; Markov chain methods for analysis of longevity, heterogeneity, and individual stochasticity; reactivity and amplification analyses). The introduction of new methods is not slowing down; if anything it is accelerating.

The COMADRE Animal Matrix Database was founded at the Max Planck Institute for Demographic Research (MPIDR) by Salguero-Gómez in 2011 (Figure 1), soon after joined by Jones, as well as a core committee, a science committee, and a team of digitizers (Supporting Information Appendix S1). The motivation for the creation of a database containing MPMs for animals was based on the success of its sister database, the COMPADRE Plant Matrix Database (Salguero-Gómez *et al.* 2015). Four years after its foundation, the COMADRE digitalization team has digitized, standardized, error-checked and supplemented information contained in over 400 species. As with the commitment for COMPADRE, more data will be released periodically (Figure 1) through the COMADRE online portal (www.comadredb.org).

## What is in the COMADRE portal?

The COMADRE portal (www.comadre-db.org) facilitates open access to the *R* data object that contains the database itself, as well as the COMADRE user’s guide. The latter contains details on the organization of the data object, the meaning and possible values for the variables within, and information on error-checks and quality controls. Additionally, Frequently Asked Questions (FAQs) can be found in the online portal (http://www.compadre-db.org/Compadre/Help).

The basic data item in COMADRE is the population projection matrix. A basic (*i.e.,* linear and time-invariant) MPM can be written

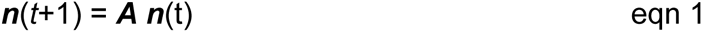

where ***n*** is a vector giving the abundance of a set of age/size/ontogenetic classes and ***A*** is a population projection matrix. The structure of the projection matrix ***A*** depends on the choice of life cycle stages and the projection interval.

In COMADRE, the projection matrix is decomposed as

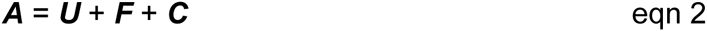

where ***U*** is the matrix describing transitions and survival of extant individuals, and ***F*** and ***C*** are the matrices describing production of new individuals by sexual and clonal reproduction, respectively. Some studies do not measure reproduction, reporting only the transition matrix ***U***. In these cases, this is reflected in the variable *MatrixFec* (see Table 1 and COMADRE User’s Guide for details). The column sums of ***U*** give the survival probabilities of the stages, and thus should not exceed 1. Some studies report **U** matrices whose column sums do exceed 1; these are noted in the database (variable *SurvivalIssue* in Table 1) and must be treated with caution.

The simple model (1) can be extended in several ways. *Seasonal* MPMs divide the year into seasons (not necessarily of the same length) and report a projection matrix ***A****_i_* for season *i*; the database entries for such seasonal models report all of the seasonal matrices. *Stochastic, density-dependent, and environment-dependent* MPMs are increasingly common in animal studies. In such cases, the MPM can be written

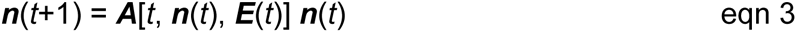

where ***E***(*t*) is some measure of environmental conditions. Such a model is associated not with a single projection matrix, but with a function that, given a time and/or environment and/or population vector, returns a projection matrix. Because such functions require a different data structure, such MPMs are not included in COMADRE 1.0.0, but we will include them in future versions.

Associated with the projection matrices is a rich set of descriptive information and metadata; thus the *R* object `COMADRE_v.1.0.0.Rdata` contains three main branches: *`metadata, matrixClass`* and `mat.` The metadata can be accessed in *R* with the command `comadre$metadata,` and it includes information about taxonomy, additional details of the study including its source, geo-location, and some details about the specific MPMs (Variables 1 through 54 in Table 1). Information about the classes used to construct the specific MPM can be accessed with the *R* command `comadre$matrixClass.` Lastly, the population projection matrices can be retrieved with the command `comadre$mat.` Data pertaining to particular matrices can thus be obtained using R’s data indexing facilities i.e. `comadre$metadata[n,]` and `comadre$matrixClass[[n]]` will return the metadata and class information pertaining to the *n*th matrix `(comadre$mat[[n]]).`

In some cases, the original data source provided information that allowed us to split the full life-cycle matrix *(matA)* into survival-dependent processes *(matU),* sexual reproduction *(matF),* and clonal reproduction, *(matC)* as described in equation (2); see variable *MatrixSplit* in Table 1. These matrices can be obtained with ease: `comadre$mat[[n]]$matA, comadre$mat[[n]]$matU etc.` Splitting the matrices in this way allows for faster, semi-automatic calculation of demographic output on hundreds of records in a few seconds. The unique matrix-specific indices allow the user to relate the information contained in the three branches of COMADRE with just a few lines of code (see examples in our GitHub repository, linked through Supplementary Online Material S3).

## COMADRE and COMPADRE: similarities

The core data in both COMADRE and COMPADRE are the population projection matrices that make up MPMs. A comparison of Table 1 in this manuscript and Table 1 in the introduction to COMPADRE (Salguero-Gómez *et al.* 2015) reveals a number of similarities. Moreover, the data quality controls are the same for COMADRE and for COMPADRE. These were detailed in an earlier publication (Salguero-Gómez et al. 2015). Due to its importance, however, we emphasize the variable *SurvivalIssue* (Table 1). The stage-specific survival of any column sums of *matU* must be a value between 0 and 1. Values greater than 1 render most analyses of survival and longevity impossible. When probabilities exceeded the error margin for rounding error and were considerably greater than 1, authors were contacted for clarification. In some cases (<13% of MPMs with this issue), these personal communications have resulted in amendments from the originally published matrices, or in the re-assignment of proportions of each matrix element in *matA* to the submatrices *matU, matF* and *matC* (Table 1). MPMs with this concern are periodically checked and, when necessary, additional clarification is requested from the authors and stored in the variable “Observation” (Table 1). Currently, only 1.2% of the MPMs (19 out of the 1,575) in version 1.0.0 have at least one life stage with survival >1.

## COMADRE and COMPADRE: differences

In spite of the similarities, animals pose some important differences that cannot be fully accommodated by the database framework of COMPADRE. The following there are key differences between the two databases:

- The variables *GrowthType, DicotMonocot* and *AngioGymno,* which are specific to plants, naturally do not exist in COMADRE.
- The variable *TPLVersion,* which identifies the taxonomic validity of plant names from The Plant List (http://www.theplantlist.org), is here substituted with CoLCheckOK, its analog for the animal kingdom via The Catalogue of Life (CoL) (http://www.catalogueoflife.org). This variable simply indicates whether or not (TRUE/FALSE) the taxonomy given in COMADRE has been validated at CoL. In just three cases the species were not present in CoL. In all other cases the taxonomic placement has been validated as correct.
- We have added the variable *MatrixFec,* which indicates whether the reproductive component (matrices ***F*** and/or ***C***) of the matrix model is missing or not. Users are cautioned to carefully examine models in which the reproductive components are missing before using them for any demographic analyses that require the full life cycle (e.g. population growth rates *λ* and its elasticities/sensitivities, damping ratio *ρ*, etc.). However, other metrics are still valid in these models (e.g. life expectancy from *matU;* table 1). *MatrixFec* will also be added to the next version of COMPADRE.
- Unlike in COMPADRE, where we reconstructed a phylogeny for plant species, a phylogeny for most animal species in COMADRE has been recently published (Hedges *et al.* 2015). Furthermore, species-level resolved trees also exist for some taxonomic groups such as mammals (Bininda-Emonds *et al.* 2007), birds (Jetz *et al.* 2012), or reptiles (Pyron, Burbrink & Wiens 2013).

## Scope and coverage of COMADRE

The current version in the COMADRE portal contains an unprecedented sample size for information on animal population dynamics: 1,575 MPMs from 272 studies corresponding to 402 species according to the authors, or 349 accepted taxonomically according to the Catalogue of Life. This represents a substantial improvement in sample size and ancillary information (Table 1) considered to date, including important comparative works examining various aspects of life history traits and population dynamics of mammals (50 species in Heppel, Caswell and Crowder 2000), birds (49 species in Sæther & Backer 2000), fish (88 species in Vélez-Espino, Fox & McLaughlin 2006), order Carnivora (285 species in van de Kerk *et al.* 2013) and various animal taxa (17 species in Jones *et al.* 2014). It must also be noted that the information analysed in those studies was not made publically available, although M. van de Kerk kindly provided COMADRE with her MPMs.

COMADRE offers a broad geographic coverage of animal population dynamics (Figure 2.A). Information in COMADRE 1.0.0 includes MPMs from all continents except Antarctica – although MPMs for Antarctic species do exist and will be released in future version of COMADRE (e.g., Emperor penguin, Jenouvrier *et al.* 2012; Antarctic petrel, Descamps *et al.* 2015). Importantly, geographic gaps do exist in our knowledge of animal demography in certain regions, including Oceania (8.13% of MPMs), and Asia (2.3%; Figure 2.B). Together, the USA (31.7%), Canada (8.8%), Australia (5.2%), and Kenya (4.8%) comprise over half the MPMs in COMADRE 1.0.0, and a clear bias exists towards terrestrial studies at low elevations (Figure 2.C). Unfortunately, few studies report MPMs from biodiversity hotspots such as Honduras, Guatemala, the Democratic Republic of Congo, Paraguay, India, and Indonesia. Furthermore, even some developed countries, such as Saudi Arabia, Italy, Greece, Ireland, Brazil and France, are under-represented.

Individual and seasonal population projection matrices (Table 1 #42) together total over 50% (783) of the projection matrices in COMADRE 1.0.0, representing unique combinations of studies × species × populations × treatments × periods (Figure 3.A). The remaining 755 projection matrices are element-by-element arithmetic means of other matrices, or constructed based on data from multiple sources (“pooled”). Given the intra-annual (Figure 4.A), inter-annual (Figure 4.C) and spatial replication (Figure 4.B) in many studies, the high proportion of mean and pooled matrices suggests a tendency in animal demographic studies to publish only summary MPMs. We encourage authors to do so as part of the supplementary materials for their papers. Authors willing to share additional matrices to be archived in COMADRE can do so by submitting the materials at.

**Figure 3.**
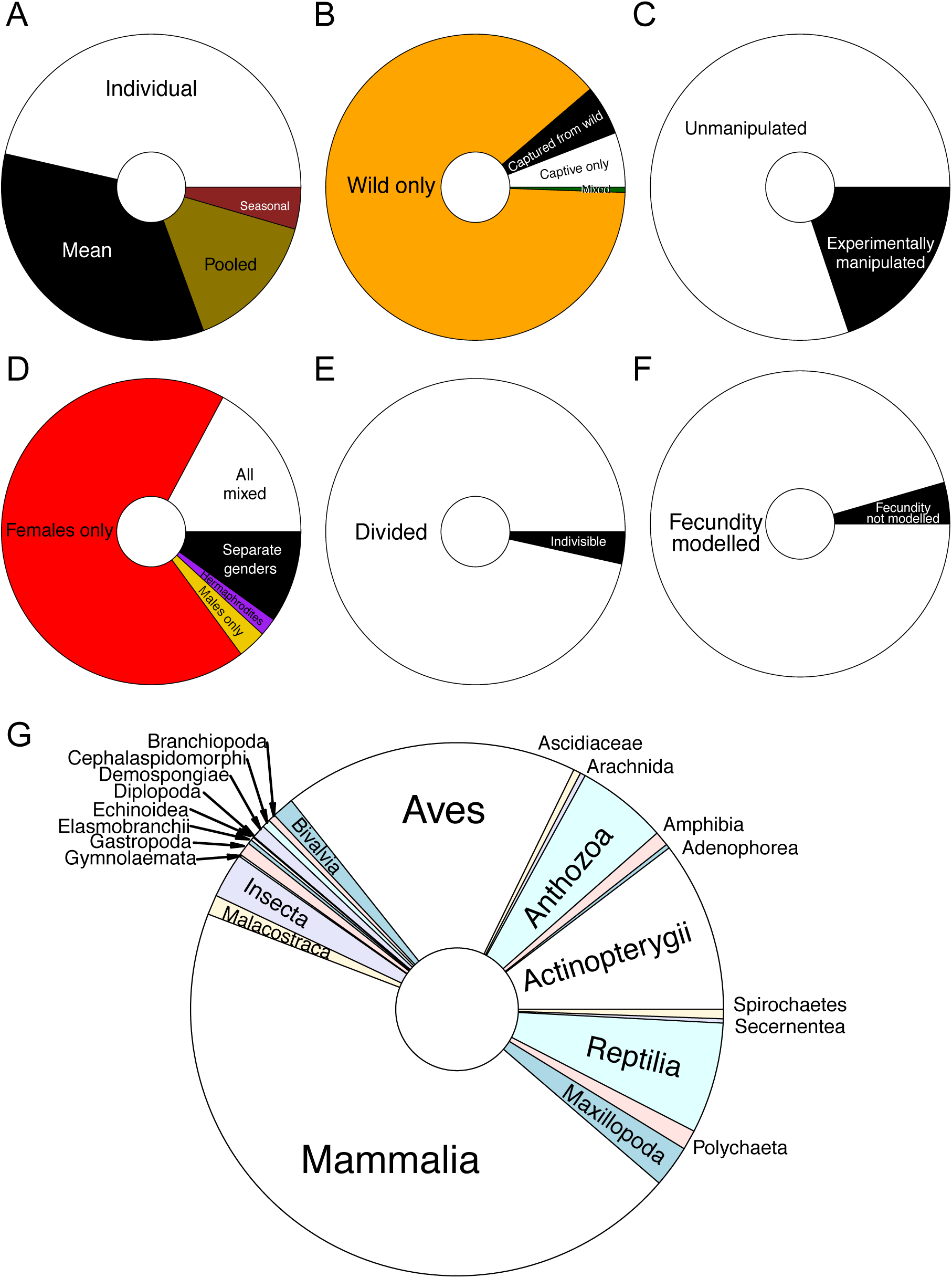
Classification of Matrix Population Models (MPMs) in COMADRE. **A.** By type of matrix (See *MatrixComposite* in Table 1). **B**. By environmental conditions of studied population (*Captivity*). **C.** By general type of treatment of matrix model under consideration (*MatrixTreatment*). **D.** By whether sex was modelled (*StudySex*). **E.** By whether the matrix ***A*** was split into submatrices ***U, F*** and ***C*** (*MatrixSplit*). **F.** By whether reproduction was modelled (*MatrixFec*). **G.** By taxonomic class representation (*Class*).

**Figure 4.**
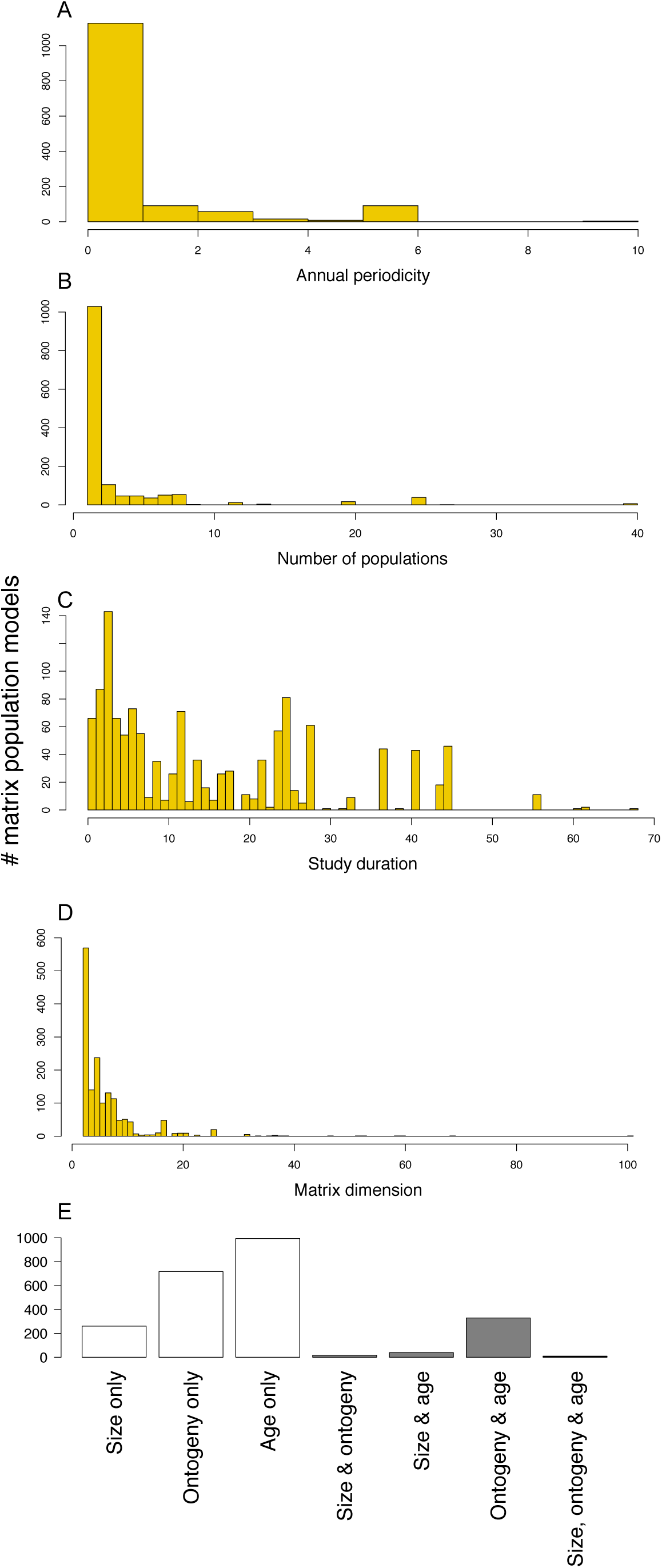
General aspects of tempo-spatial replication and stage construction of matrices in COMADRE. **A.** Frequency of matrix models by demographic census periodicity on an annual basis (See *Periodicity* in Table 1). **B.** Number of populations *(NumberPopulations).* **C.** Duration of the study (in years; *StudyDuration*). **D.** Matrix dimensionality *(MatrixDimension).* **E.** Criteria used to construct the matrix model *(MatrixCriteriaSize, MatrixCriteriaOntogeny* & *MatrixCriteriaAge*).

Most studies in COMADRE are of natural populations in the wild (87%, Figure 3.B), and under unmanipulated conditions (80%; Figure 3.C). Most of the demographic studies in COMADRE are based on females only (68%; Figure 3.D); this is common practice in animal demographic studies (particularly in mammals), since quantifying reproduction is usually easier in females than in males. We have noted, in the variable “Observations” (Table 1 #52), when the primary sex ratio was stated by the author as differing from 1:1 (female:male). For the vast majority of matrices (97%; Figure 3.E), we have successfully split the full matrix ***A*** into its subcomponents of survival (***U***), sexual reproduction (***F***) and clonal reproduction (***C***), and only 4% of the MPMs do not incorporate reproductive information (Figure 3.F).

The data in COMADRE 1.0.0 represent a wide range of animal groups (Figure 3.G). However, there are some strong taxonomic biases. Mammals represent 44.4% of the MPMs in the current version of COMADRE, followed by birds (17.9%), bony fish (10.3%), and reptiles (6.7%). We include very few MPMs for amphibians, despite global concerns for their conservation status (Beebee & Griffiths 2005; Wake & Vredenburg 2008) or for insects (2.71%), despite their high species richness, estimated to comprise the majority of the animal kingdom (Hedges *et al.* 2015). The latter is particularly surprising since the early developments of MPMs focused on insects thanks to their clearly structured population dynamics (Lefkovitch 1965; Rabinovich 1969). Aside from bony fish (Actinopterygii), we also lack significant amounts of demographic information on marine organisms in COMADRE, including corals (5.5%), bivalves (1.33%), sponges (0.8%), sea urchins (0.12%), and cartilaginous fish (0.2%). No information in COMADRE 1.0.0 exists for the infraclasses Marsupialia (kangaroos, wallabies, koala, possums, opossums, wombats, etc.) or order Struthioniformes (kiwis, emu, ostriches, etc.), neither in the additional 800 species records that are currently being digitized and error-checked for future version releases.

The replication of studies through time and space is highly variable in COMADRE. Yet, the average duration of studies in COMADRE 1.0.0 (15.69 years ± 13.80 S.D.; Figure 4.C) is greater than in plant MPM studies (Salguero-Gómez et al. 2015). Long duration is essential for many demographic studies in the animal kingdom, as some animals, such as the clam *Arctica islandica,* giant tortoises *(Geochelone nigra, G. gigantea),* rockfish (*Sebastes* sp.), and the bowhead whale *(Balaena mysticetus),* can reach over 150 years of age (de Magalhaes & Costa 2009). Notable demographic studies using MPMs parameterized with long time-series include *Vipera aspis* (17 years, Altwegg *et al.* 2005), *Ursus americanus* (22 years, Mitchell *et al.* 2009), *Delphinus delphis* (35 years, Mannocci *et al.* 2012), *Recurvirostra avosetta* (40 years, Hill 1988), *Elatobium abietinum* (41 years, Estay *et al.* 2012), *Marmota flaviventris* (44 years, Ozgul *et al.* 2009), *Haliaeetus albicilla* (62 years, Krüger, Grünkorn & Struwe-Juhl 2010), *Diomedea exulans* (51 years, Barbraud *et al.* 2013), and *Aythya affinis* (72 years, Koons *et al.* 2006).

In contrast to the duration, the average number of populations considered in each study is low, averaging 3.31 ± 5.26 (S.D.). The low spatial replication currently limits much-needed understanding of the geographic variability of demographic rates within species. Recently initiated efforts to increase spatial replication for certain species in the plant kingdom (e.g. on *Plantago lanceolata,* PlantPopNet, www.plantago.plantpopnet.com) are a useful model that could be replicated in future work on animals. It is perhaps not surprising that the animal studies with highest spatial replication in COMADRE 1.0.0 focus on humans, with the foundational archive of human MPMs compiled by Keyfitz and Flieger (1968), which covers populations from 156 countries. We note that analyses of spatial and other kinds of variability in animal population studies are becoming more sophisticated due to the use of model selection methods to explicitly include environmental variables (e.g., Thomson, Cooch & Conroy 2009), and the concept of “spatial replication” used in plant studies may acquire a different meaning to that used in most animal studies.

## Unlocking global analyses

> *“I’m not interested in your data; I’m interested in merging your data with other data. Your data will never be as exciting as what I can merge it with”*
>
> — Tim Berners-Lee

The scientific promise of the COMADRE Animal Matrix Database does not reside exclusively in its hundreds of MPMs, but also in the many outputs that can be derived from them, and the possibility to put them in a broader spatial, ecological and evolutionary context using other open access databases and user collected or compiled data. Users of COMADRE can find several *R* scripts available to manipulate and interact with matrices, and derive basic demographic outputs (Supporting Information Appendix S3, and our growing GitHub repository). Users are welcome to explore these or other more developed open-source libraries (Stubben & Milligan 2007; Stott, Hodgson & Townley 2012; Metcalf *et al.* 2013), and to carry out their own calculations based on methods for the analyses of MPMs (e.g., Caswell 2001; Morris & Doak 2002).

The schedules of growth, survival, and reproduction and the associated population performance metrics available through COMADRE will enable further comparative analyses of life history variation and population performance relative to the environment. For example, information in COMADRE can be integrated with existing repositories for other data such as genetic sequences (GenBank; Benson et al. 2013), distribution and occurrences (GBIF; Flemons *et al.* 2007), and conservation status and threats (BirdLife, http://www.birdlife.org/datazone/; IUCN Red List, http://www.iucnredlist.org). Data on species-level life history traits are also available for specific taxonomic groups including vertebrates (AnAge; de Magalhaes & Costa 2009), mammals (Ernest 2003; PanTHERIA, Jones et al. 2009), amphibians (Trochet *et al.* 2014), fish (FishBase), and reptiles (SCALETOOL, www.scale-project.net). Lastly, an upcoming resource, DATLife (Scheuerlein *et al.* unpublished), containing age-specific mortality and ancillary data on animal species will be of particularly interest to supplement COMADRE, as methods already exist to convert life tables into MPMs and *vice versa* (Caswell 2001). In addition to this rich and rapidly growing body of data, a diverse set of tools are emerging that will facilitate these large-scale comparative analyses including the *R* packages *taxize* (Chamberlain & Szöcs 2013), *letsR* (Vilela & Villalobos, 2015) which facilitate taxonomic matching and macroecological analyses respectively.

The compilation of demographic data in COMADRE will also enable the identification of gaps in our knowledge of animal population dynamics and, as is now happening for plants, will catalyze new studies at broad spatial scales. The open-access publication of both COMPADRE and COMADRE databases will facilitate further comparative demographic analyses across plant and animal kingdoms (see Jones *et al.* 2014) enabling tests of life history and population dynamics theory across a wide range of species with contrasting life histories. We suggest that researchers revisit the canonical tenets of animal life history to confront established theories with data compilations that are vastly richer than was available 30 years ago.

While the COMADRE team strives to make the database as accurate as possible, we make no claims, promises, or guarantees about the accuracy, completeness, or adequacy of the database, and expressly disclaim liability for errors and omissions in the contents of the database. No warranty of any kind is given for any particular use of COMADRE. Users who detect apparent errors are encouraged to contact us at.

Although this first release contains 402 species, we have already identified over 800 additional animal species with MPMs, and our on-going efforts will release them as they become fully-digitized, error-checked and supplemented in the coming years (Salguero-Gómez *et al.* unpublished). Finally, researchers using the data archived here are encouraged to cite also the original sources in their works (Supporting Information Appendix S4).

We are extremely grateful to the many ecologists, zoologists, and evolutionary biologists who have made the projection matrices from their MPMs available for publication in this open access database. Some of the data stored here must rank among the most valuable (and most expensive to collect) biological information in existence. The providers of data have shared the vision of the COMADRE leaders, that important data should be made available to all interested parties for free. In return, we simply encourage demographers, conservation biologists, ecologists and evolutionary biologists worldwide to mine this database and find out as much as they can about global, phylogenetic and ecological patterns in animal life history and population dynamics.

## Acknowledgments

The original author list whose works constitute the motivation for the COMADRE Animal Matrix Database version 1.0.0 can be found in the Supporting Information Appendix S4. The authors’ willingness to share unpublished data and clarify questions on published materials is an inspiration for the new era of big, open, data. COMADRE is currently supported by the Evolutionary Demography Laboratory at the Max Planck Institute for Demographic Research (MPIDR). For a list of funding support since its inception see Supporting Information Appendix S5. Logistical and computational support was provided by the IT team of the MPIDR.

## Data accessibility statement

The data associated with this manuscript can be accessed at www.comadre-db.org

## Supporting Information

Additional Supporting Information may be found in the online version of this article.

**Appendix S1.** Constituents of COMADRE.

**Appendix S2.** COMADRE user’s guide.

**Appendix S3.** COMADRE *R* scripts.

**Appendix S4.** Extended literature used in COMADRE 1.0.0.

**Appendix S5.** Funding and extended acknowledgements.

**Appendix S6.** Author contributions.

**Appendix S7.** Supporting information references.

As a service to our authors and readers, this journal provides supporting information supplied by the authors. Such materials may be re-organized for online delivery, but are not copy-edited or typeset. Technical support issues arising from supporting information (other than missing files) should be addressed to the authors.

## Footnotes

1 A glance at Hutchinson’s (1978) population ecology text, based on a course he taught for many years at Yale, will show how influential life table methods were, and how intimately connected the approaches of animal demographers were to those of human demographers.

